# Metabolic traits and the niche of bulk soil bacteria in a Mediterranean grassland

**DOI:** 10.1101/2022.06.21.497019

**Authors:** Kateryna Zhalnina, Richard Allen White, Markus de Raad, Kai Deng, Carrie D. Nicora, Ulas Karaoz, Jennifer Pett-Ridge, Mary K. Firestone, Mary S. Lipton, Trent R. Northen, Eoin L. Brodie

## Abstract

Soil microorganisms have adapted to compete and exploit different metabolic niches in their physically and chemically diverse environment via evolution and acquisition of distinct physiological and biochemical traits. As the interface for most carbon and nutrient exchange between plants and microorganisms, the rhizosphere has received substantial attention. By comparison, what is commonly termed bulk-soil (soil free of living roots) represents a far greater volume and surface area throughout the season, and substantially higher taxonomic and phylogenetic diversity; the traits and activity of its inhabitants may also have a significant impact on overall soil function. We used a combination of comparative genomics and exoproteomics to identify metabolic traits of bacteria adapted to life in bulk soil and compared these with traits of bacteria living in the rhizosphere of wild oat, *Avena barbata*. In bulk soil bacteria, we observed: (i) greater investment in extracellular polymer-degrading enzyme production; (ii) greater potential for secretion (presence of signal peptides) of polymer-degrading enzymes; (iii) production of accessory proteins (carbohydrate binding modules) fused with glycoside hydrolases that enhance substrate affinity, stabilize, and increase reaction rates of polymer degrading enzymes; and (iv) organization of polymer degradation machinery within gene clusters that facilitate co-transcription of enzymes, transcription factors and transporters for polymer depolymerization products. Together, these findings suggest that unlike rhizosphere-adapted bacteria—which specialize on small molecules released primarily as root exudates—bulk soil-adapted bacteria have evolved to exploit plant polymers. This biochemically costly strategy may be mitigated by protein-level adaptations that enhance the efficiency of extracellular enzyme-mediated substrate acquisition.

**IMPORTANCE:** Plant-soil-microbe interactions are dynamic and complex, with significant implications for ecosystem functioning. Microbial traits, such as nutrient acquisition and growth yield, combined with soil and climate parameters, impact major biogeochemical processes and can define the future fate of soil carbon. Diverse soil microorganisms occupy different physical habitats within soil and exploit distinct niches by expressing different metabolic traits. Identifying and quantifying traits that underlie their fitness and function is key for understanding and predicting how soil carbon transformation and stabilization will change in the future or can be managed through intervention.

## Introduction

Soils contain a wide variety of niches due to their physical and chemical heterogeneity and it is increasingly apparent that this niche complexity underlies the nature of soil as a hotspot for microbial diversity. Microorganisms that compete and survive in soils are adapted to a wide array of niches with distinct chemical forms, concentrations and availability of carbon and nutrients provided in different physical habitats. Based on macro-ecological theory, we would expect that life history traits and specific habitat adaptations should be evident in the combination of physiological, genomic and biochemical traits reflected by soil microorganisms gathered from different soil locations (1). There is a particularly compelling imperative to derive genotype to phenotype relationships to begin to generalize the suite of traits required to subsist and succeed in the varied niches of the soil environment. However, observations and quantification of these traits are currently lacking.

Rhizosphere and bulk (soil free of living roots) soil represent distinct chemical environments even in situations when they share similar mineralogical and physical properties (2, 3). Plant growth drives a bloom of microbial activity and biomass growth near developing roots due to the release of a significant fraction of photosynthate as exudates. By contrast, soil from outside the zone of living root (>4 mm from living root) influence represents a less variable environment with lower resource availability and proportionally higher importance of polymeric carbon substrates(4–7). Due to high density of roots in the grasslands detritusphere and bulk soil represent the same living environment characterized by the input of recalcitrant highly polymeric organics from litter upon root death at various depths (8, 9). The contrasting resource availability between rhizosphere and bulk soils, including detritusphere, encourages growth of microorganisms with distinct properties, though both are drawn from a common seed bank of microorganisms (4, 10). While the genetic and metabolic properties underlying traits required for successful rhizosphere colonization have been extensively explored (11–19), the traits and underlying mechanisms of organisms adapted to bulk soil have received significantly less attention (20, 21).

What defines bulk soil microorganisms and shapes their broader impacts on soil function? Decomposition of polymers in bulk soil represents one of the central spots of microbial activities or “hot spots” in soil (8). In contrast to the much higher but short-lived microbial activities in the rhizosphere, bulk soil microbial communities have a larger overall volume and prolonged activity, and significantly influence soil C dynamics and stabilization (8, 22). Additionally, bulk soils have substantially higher taxonomic and phylogenetic diversity (4, 10, 18), a higher proportion of microorganisms adapted to oligotrophic conditions (10), and higher capacity for macromolecular carbon degradation (20). Our previous investigation of soil isolates substrate preferences suggests that while all soil bacteria tested could use common metabolites released by roots (e.g. glucose or adenine;(11)), bulk soil bacteria had a greater genomic investment in the use of plant polymers, particularly the capacity to produce enzymes classified as glycoside hydrolases (GHs).

Glycoside hydrolases are the most abundant of the carbohydrate active enzymes (CAZymes) and many participate in the deconstruction of complex carbohydrates via hydrolysis of glycosidic bonds. Microbial GHs are widely distributed across different environments, including soil, marine, phyllosphere, human skin and gut (23) and insect digestive tracts (24). GH content of genomes can be extremely variable across microbial phyla and indeed between bacterial strains (25) (26–28). In soil, stimulation of GH-containing microorganisms has been invoked to explain organic matter priming (20, 29, 30). Frequently, carbohydrate binding modules (CBMs) work together with GHs to enhance hydrolysis of insoluble polymers and increase depolymerization efficacy of the enzymes (31). However, while rhizosphere taxa appear to prefer some low molecular weight compounds (11, 20), both rhizosphere and bulk soil bacteria carry a large variety of CAZy enzymes in their genomes and it is not clear whether bacteria that are considered bulk soil-adapted have a competitive advantage to deconstruct root-derived polymers.

Extracellular enzymes secretion by bacteria is associated with microbial strategy for nutrient acquisition in soils via depolymerization of complex substrates, such as cellulose, hemicellulose, starch, chitin, protein and lignin (32–36). As the act of enzyme secretion for resource acquisition is a yield trade-off for such organisms (37), specialization within this niche may come at the cost of reduced growth yield(38). Given the potentially high costs of protein secretion, selection for more efficient use of secreted proteins might also be expected. Selection for protein-level traits, for example carbohydrate binding modules (CBMs) that increase the affinity of enzymes for substrates, enhance processivity, and reduce advective or diffusive loss of enzymes from solid polymeric substrates, could offset costs by enhancing efficiency.

Here, we asked the following: (1) What are the key metabolic traits of bulk-soil bacteria and do these traits confer a fitness advantage? and (2) is there evidence of metabolic evolutionary strategies that may reduce costs of extracellular enzyme production by enhancing efficiency of these enzymes. To answer these questions, we analyzed the distribution of CAZy enzymes in the genomes of 28 bacterial isolates identified as either rhizosphere- or bulk soil-adapted in previous studies of *Avena barbata* (10, 11). We explored differences in extracellular proteins secreted by both groups of bacteria, and used exoproteomics to evaluate enzymes produced and released by a subset of the isolates grown with four substrate treatments: monomer-rich medium (to mimic the actively exuding plant root), or media with polymeric substrates (to mimic detritus found in bulk soil). Using Matrix Assisted Laser Desorption Ionization Mass-Spectrometry (MALDI-MS), we also measured activities of polymer degrading enzymes for isolates grown on these four substrate treatments. Our results provide evidence for substrate preference-based niche differentiation and linked organism-level and protein-level traits associated with life in the bulk soil environment.

## RESULTS

### Genome distribution of CAZy enzymes and GH associated carbohydrate binding modules

To analyze the genomic potential of soil microbes to use plant polymers, we selected 28 genome-sequenced isolates (Table S1) that represent abundant and taxonomically diverse bacterial taxa in a Northern California Mediterranean grassland soil and that have been enriched either in rhizosphere or in bulk soil environments (11). Previously we isolated these bacteria from this grassland soil and studied the response of these isolates to growth of the common annual grass *Avena barbata*. We mapped the isolates’ 16S ribosomal RNA genes to the total OTUs detected during *Avena* root development{Shi:2015gx(10) and classified them into two response groups (11). The first group includes isolates mapped to the OTUs that increased in relative abundance in response to plant root growth (termed ‘rhizosphere bacteria’ herein), while the second group includes isolates that were more abundant in the bulk soil and declined in relative abundance during plant growth (termed ‘bulk soil bacteria’) (Fig. S1 and Table S1).

We analyzed the distribution of carbohydrate active enzyme (CAZy) families in the genomes of these two groups of bacteria (Table S1 and Fig. S2). Genomes of bulk soil bacteria contained significantly more total CAZy genes (*P*=0.004) as well as more specific families, such as carbohydrate esterases (CE), glycoside hydrolases (GHs), carbohydrate binding modules (CBMs) compared to the rhizosphere bacteria (Fig. 1A). Frequently CBMs work together with GHs to enchance hydrolysis of insoluble polymers and increase depolymerization efficacy of the enzymes (31). We identified that bulk soil bacteria had higher numbers of fused GH-CBM domains when accounting for genome size and a higher overall percentage of fused GH-CBM domains across all GHs (Fig. 1B). In addition we found that bulk soil bacteria had higher number of GHs and percentage of total GHs organized in the clusters, consisting of at least one CAZyme, a transporter and a transcription factor gene (Fig. 1B).

**FIG 1.**
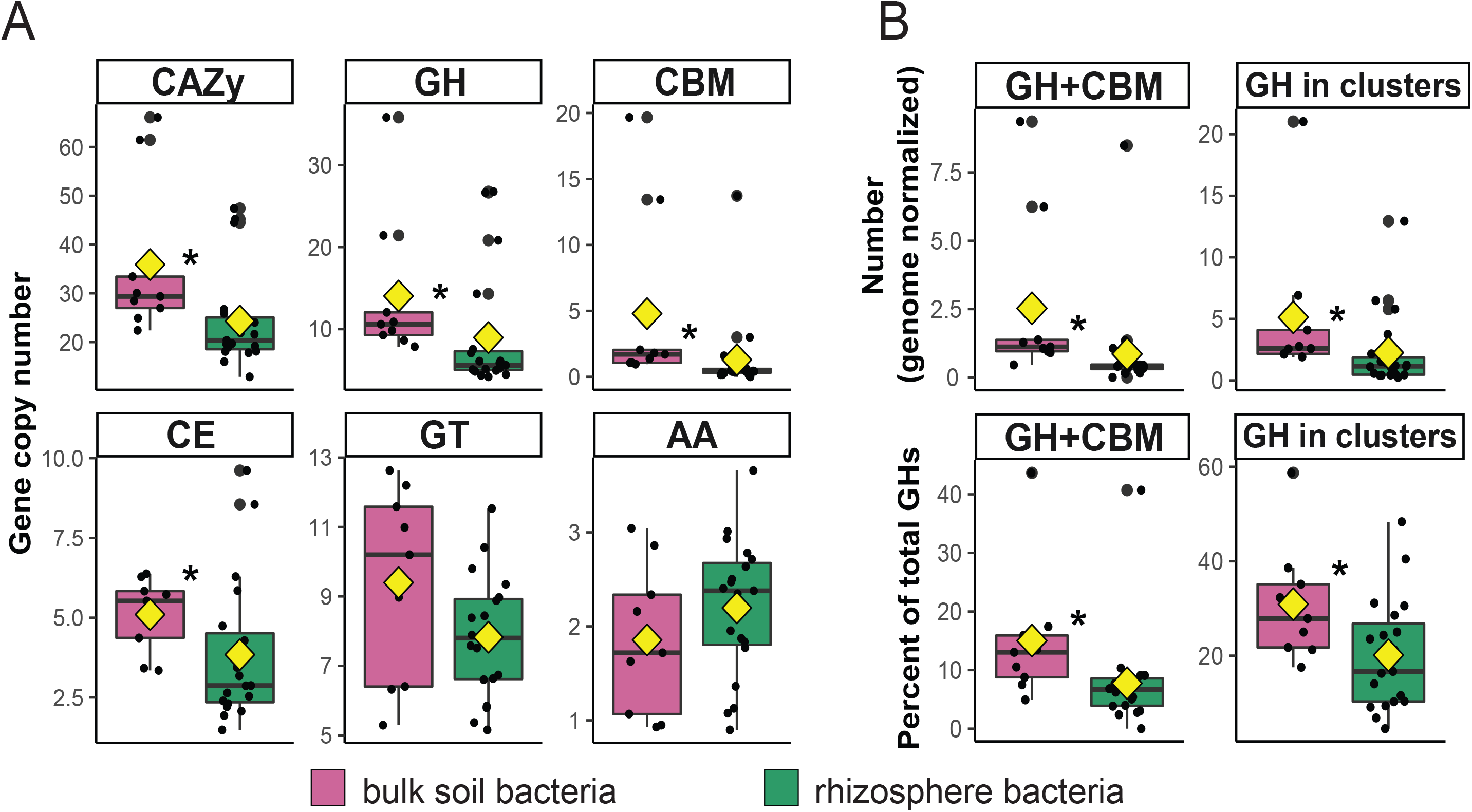
CAZy families, fused domains and gene clusters in the genomes of rhizosphere and bulk soil bacteria. (**A**) Carbohydrate active enzyme (CAZy) families in the genomes of 28 isolates from a California annual grassland soil. These isolates were classified into two groups based on their abundances in rhizosphere of the common annual grass *Avena barbata* and bulk soil (10, 11): i) bulk soil bacteria (n=9), which were more abundant in bulk soil and declined in relative abundance in response to the plant growth, (ii) rhizosphere bacteria (n=19), which were more abundant in the rhizosphere. AA – auxiliary activities enzymes; CE – carbohydrate esterases; GT – glycosyl transferases; GH – glycoside hydrolases; CBM – carbohydrate binding modules. Gene copy number was normalized by the genome size of each isolate. (**B**) Presence of fused GH and CBM domains and gene clusters containing GHs in the genomes of bulk soil and rhizosphere isolates per the genome size and per number of total GHs. Differences in the distributions of traits between the two groups of isolates (rhizosphere and bulk soil bacteria) were evaluated using a Kruskal–Wallis one-way analysis of variance test (\**P*<0.05). In each box plot, a point denotes a number of genes or fused domains of CAZy families calculated for each microbial isolate used in this study. The top and bottom of the box represent the 25^th^ and 75^th^ percentiles, the horizontal line inside each box represents the 50^th^ percentile/median and the whiskers represent the range of the points excluding outliers. Outliers are denoted as large points outside whiskers. Diamond symbols in each box plot represent the mean.

### Genomic potential for cellulose, hemicellulose and oligosaccharide degradation

Cellulose and hemicellulose are the most abundant plant polymers in soil. Across all analyzed bacterial genomes, we identified GHs that are known from the literature to target cellulose, hemicellulose and oligosaccharides (Table S1 and Fig. 2) (25). Bulk soil bacteria possessed significantly more GH families linked to the decomposition of hemicellulose and oligosaccharides than rhizosphere bacteria (Fig. 2A). Both groups have a similar number of GHs associated with cellulose decomposition. Hemicellulose degradation GH families that were found at significantly higher levels in bulk soil bacteria belonged to the CAZy families β-galactosidase/β-mannosidase (GH2), β-1,3-xylanase/β-mannanase (GH26), α-galactosidase (GH36) and β-xylosidase (GH43).

**FIG 2.**
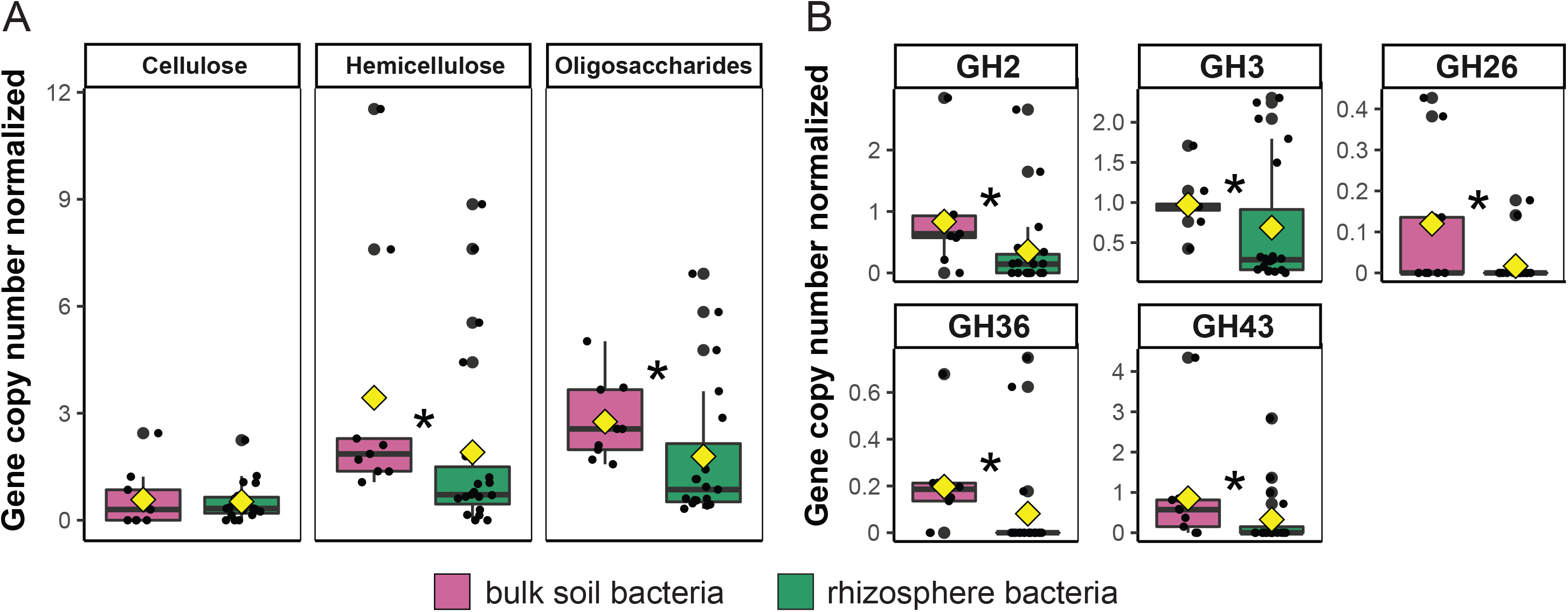
Glycoside hydrolases (GHs) in the genomes of bulk soil and rhizosphere bacteria and their target substrates. (**A**) Number of GHs known to hydrolyze cellulose, hemicellulose and oligosaccharides. (**B**) Copy number of specific GH families that target hemicellulose or oligosaccharides (n=28 genomes). All GH copy numbers are normalized by genome size for each isolate. Differences in the number of GHs between the two groups of bacteria (bulk soil and rhizosphere bacteria) were evaluated using the Kruskal–Wallis one-way analysis of variance (\**P*<0.05). In each box plot, a point denotes a number of GH genes calculated for each microbial isolate used in this study. The top and bottom of the box represent the 25^th^ and 75^th^ percentiles, the horizontal line inside each box represents the 50^th^ percentile/median and the whiskers represent the range of the points excluding outliers. Outliers are denoted as large points outside whiskers. Diamond symbols in each box plot represent the mean.

### Isolate metabolic activity on monomer-rich and plant polymer media

To determine whether the genome-predicted capacity to use plant polymers was reflected in enhanced metabolic and enzymatic activity on specific carbon (C) substrates, we selected eight bacteria isolates that represent bulk soil bacteria (n=4) and rhizosphere bacteria (n=4) (Table S1). We grew these bacterial isolates on four different substrate sources, including: (i) a 10x-diluted monomer-rich complex medium (R2A) containing simple C molecules, including sugars, carboxylic acids, nucleobases, amino acids (e.g. glucose, pyruvate, adenine, proline) (39, 40), and (ii) defined medium (M9) supplemented with ground and sterilized *Avena* root litter, or the individual polymers (iii) xylan and (iv) cellulose. To assess metabolic and enzymatic activity on these four substrate groups, we quantified ATP production via a bioluminescence assay, and measured the activity of polymer-degrading enzymes (β-glucosidase and β-xylosidase) with mass spectrometry (Fig. 3). After 20 days of bacterial growth, ATP production of bulk soil bacteria was on average 4.5 (xylan medium), 6.5 (monomer-rich), 17 (root litter) and 18 (cellulose) fold higher relative to rhizosphere bacteria (Fig. 3A). Similarly, higher glucosidase and xylosidase activities of bulk soil bacteria were observed compared to rhizosphere bacteria (Fig. 3B).

**FIG 3.**
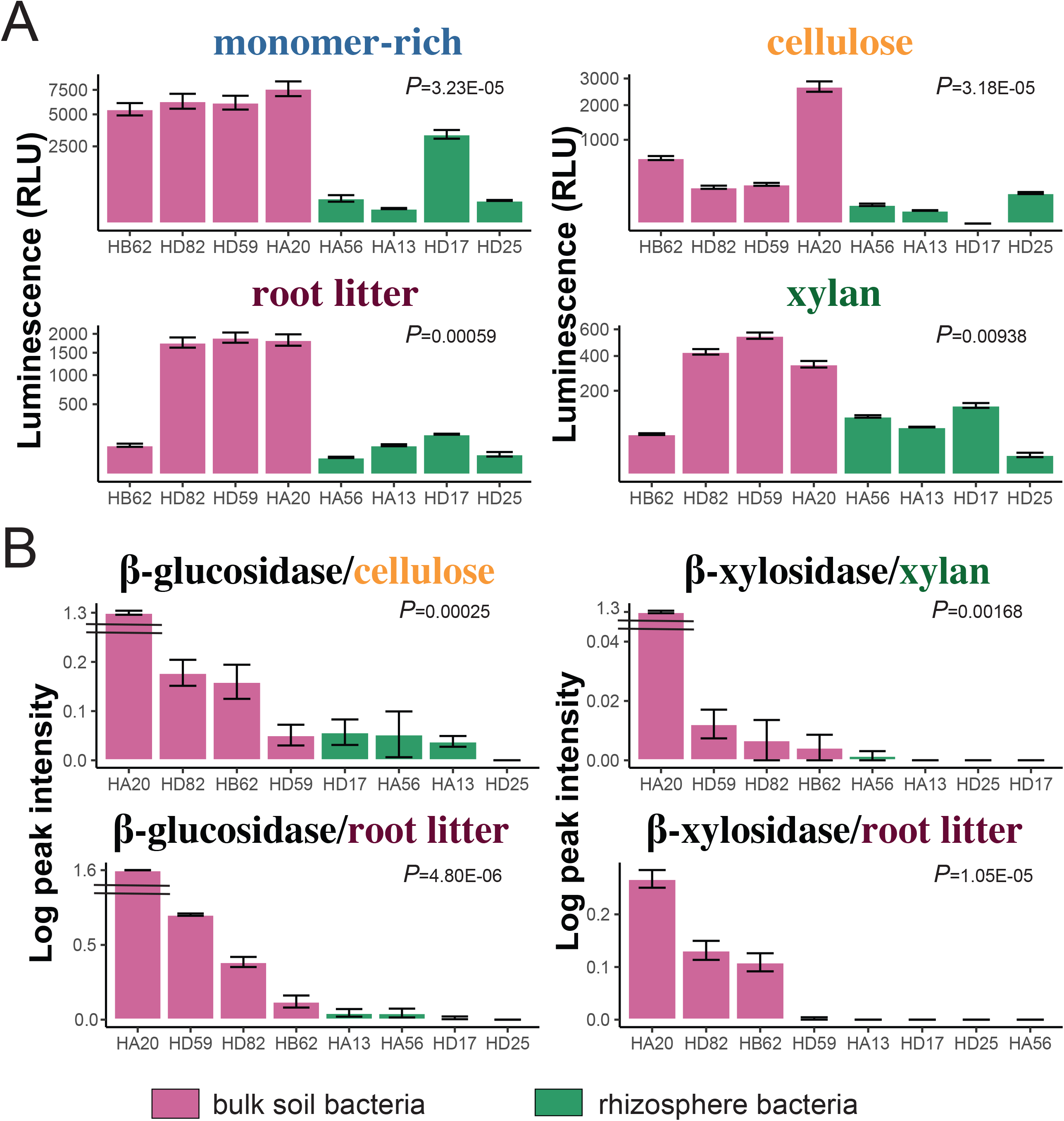
Metabolic activity of bulk soil and rhizosphere bacteria grown on different substrates. (**A**) ATP-bioluminescence of bulk soil and rhizosphere bacterial isolates after 20 days of incubation on a 10x-diluted monomer-rich medium or minimal media amended with root litter, xylan, or cellulose (n=3). RLU are relative light units. (**B**) The activity of two polymer-degrading enzymes (β-glucosidase and β-xylosidase) measured with Matrix Assisted Laser Desorption/Ionization Mass-Spectrometry (n=4). Error bars represent standard error. Rhizosphere bacteria did not exhibit xylosidase activity. Differences in ATP-bioluminescence and enzymatic activity between the two groups of bacteria (bulk soil and rhizosphere bacteria) were evaluated using the Kruskal–Wallis test.

### Extracellular enzymes produced by bacterial isolates growing on plant polymers and monomer-rich media

Using mass spectrometry-based proteomics, we analyzed extracellular proteins produced by the eight bacterial isolates growing on the four carbon substrates described above. The relative abundance of CAZy, GHs, and CBMs, is represented as a percentage of the total extracellular peptide counts, identified in each isolate replicate on each medium type in 0.1μg/μl of analyzed peptides (Table S2). Bulk soil bacteria secreted proportionally more CAZy and GH enzymes when grown on media containing plant polymers relative to the monomer-rich medium (Fig. 4). In contrast, rhizosphere bacteria secreted similar relative abundances of CAZy, GHs and CBMs on the monomer-rich medium and on the plant polymer media (Fig. 4). Rhizosphere bacteria demonstrated larger number the total extracellular peptide counts on monomer-rich medium and on cellulose than by bulk soil bacteria (Table S2). However, CAZy, GH, and CBM enzymes secreted by the rhizosphere group represented a significantly lower proportion (relative abundance) of the total extracellular peptide counts than observed for the bulk soil bacteria, regardless of substrate (Fig. 4).

**FIG 4.**
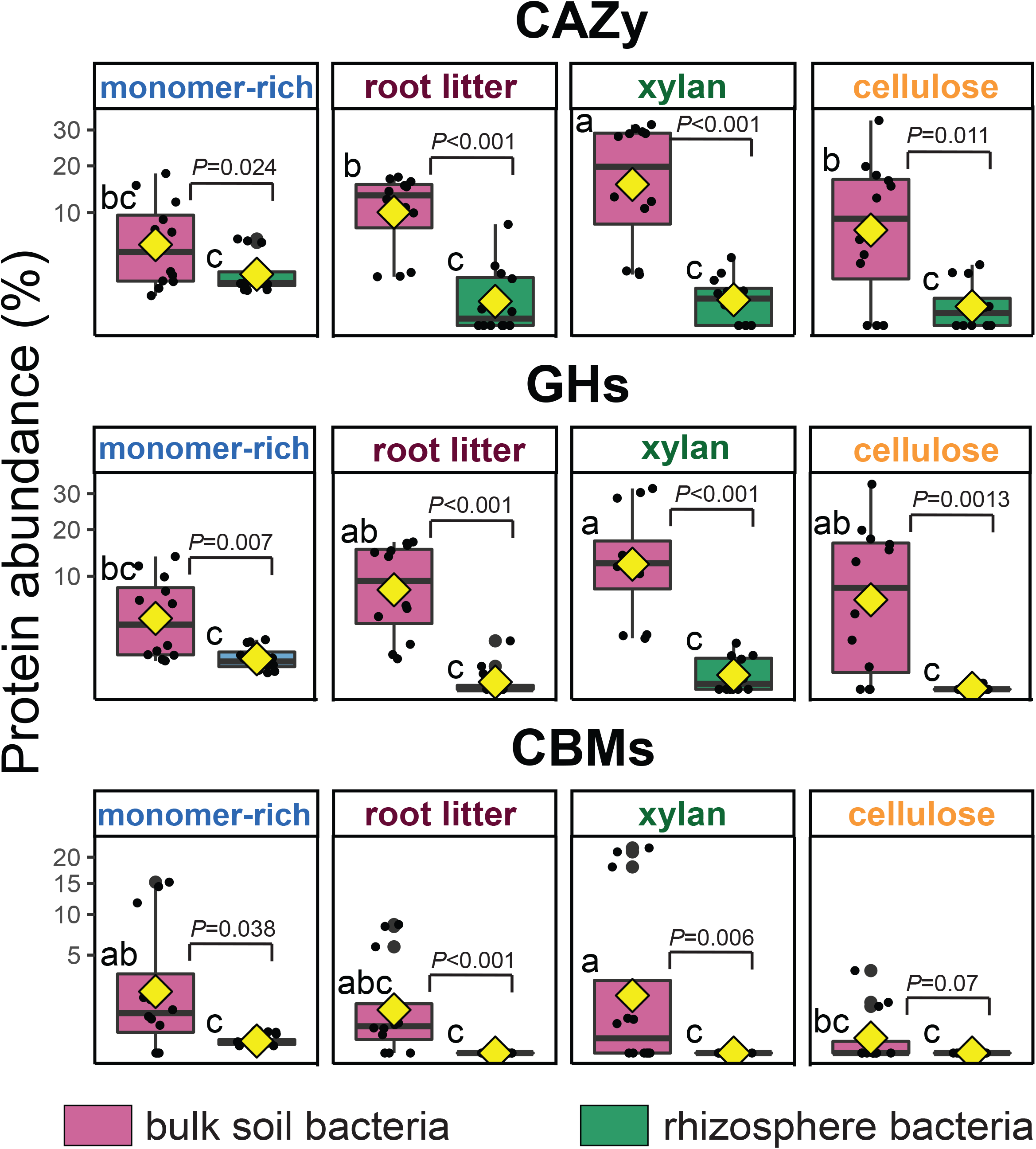
Relative abundance of carbohydrate active proteins produced by bulk soil and rhizosphere bacteria on four different media. Relative abundance of CAZy, GHs and CBMs in bulk soil bacteria and in rhizosphere bacteria, represented as a percent of the total peptide counts identified in each sample in 0.1μg/μl of analyzed peptides. Differences in the percentages of the peptide counts between bulk soil and rhizosphere bacteria on different media were evaluated using the Kruskal–Wallis test. Statistical differences across four media in bulk soil and in rhizosphere bacteria were determined by Duncan’s multiple range test. Treatments indicated by different letters are significantly different (*P*<0.05).

Within the bulk soil bacteria, we observed different strategies of plant polymer utilization. Some bulk soil bacteria appeared to secrete GHs on both xylan and cellulose (HA20, HD59), while other bulk soil bacteria produced GHs in a medium-dependent manner (e.g. GHs produced by HD82 and HB62 on xylan medium) (Fig. S4). Interestingly, although *Conexibacter* HD59 and *Solirubrobacter* HD82 belong to the same bacterial order Solirubrobacterales, and have an average nucleotide identity (ANI) just over 77%, HD59 produced extracellular GHs on both xylan and cellulose, while HD82 only secreted hydrolases on xylan – suggesting substrate specialization is poorly phylogenetically conserved (Fig. S4).

### Abundance, richness and putative activities of CBMs and GHs families secreted by bulk soil and rhizosphere bacteria

Microbial deconstruction of plant polymers includes synergistic activity of many different classes of GHs and their accessories, such as CBMs. A total of 37 GHs and CBMs families have been identified in this study (Table S3). We analyzed extracellular proteins for the presence of specific GH and CBM families produced by bulk soil and rhizosphere bacteria in response to plant polymers or monomer-rich media (Fig. 5). Both groups of bacteria had a variety of GHs and CBMs in their extracellular proteome (Fig. 5A, B). Bulk soil bacteria secreted not only a greater relative abundance (Fig. 5A, B, note distinct Y axis scales) but also a greater richness of CBMs and GHs families (total unique GHs and CBMs families n=29) than rhizosphere bacteria (total unique GHs and CBMs families n=18), particularly on plant polymer media (Fig. S5 and Table S3). A larger richness of GH and CBM families was released by rhizosphere bacteria on monomer-rich media compared to plant polymer media (Fig. 5B and Fig. S5). Essentially no GHs and CBMs were secreted by the rhizosphere isolates grown with cellulose.

**FIG 5.**
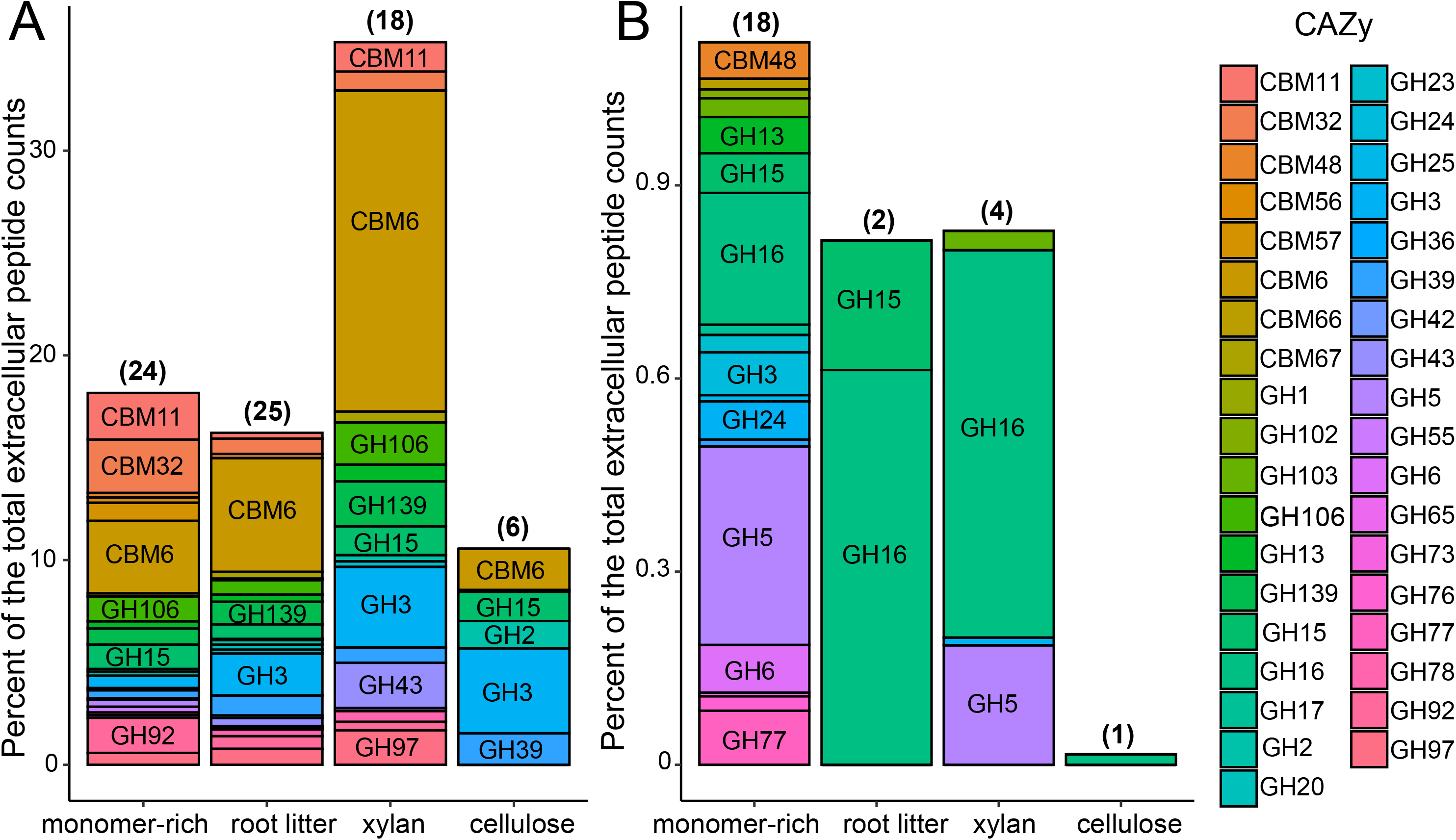
GHs and CBMs families produced on different media by the (A) bulk soil bacteria and (B) rhizosphere bacteria. Relative abundance of each GH or CBM family represented as a percent of the total number of peptide counts identified in each exoproteome sample and averaged within the enzymes of the same CAZy family for bulk soil and rhizosphere bacteria. The most abundant protein families identified are indicated on each bar (for bulk soil bacteria families with abundance >1% and rhizosphere bacteria >0.05%). Note distinct Y axis scales. Numbers in brackets represent total unique GHs and CBMs families identified across four media in bulk soil and rhizosphere bacteria.

Seven CAZy families (GH2, GH3, GH13, GH15, GH16, GH39) with β-galactosidase, β-glucosidase, α-amylase, glucoamylase, β-glucanase activities and CBM6 that binds on β-1,4-xylan or amorphous cellulose were secreted by bulk soil and rhizosphere bacteria on all four media (Fig. 5, Table S3). Twelve families (GH5, GH23, GH55, GH78, GH92, GH97, GH106, GH139, CBM11, CBM32, CBM48, CBM67) were identified on three media, excluding cellulose, and mainly secreted by bulk soil bacteria (e.g., endo-β-1,4-glucanase, chitinase, α-L-rhamnosidase, α-mannosidase, α-glucosidase, α-2-O-Me-L-fucosidase, CBMs that bind on β-1,4-glucan, galactose, glycogen). Eleven GHs and CBMs families (GH6, GH17, GH24, GH25, GH36, GH65, GH73, GH102, CBM56, CBM57, CBM66) were secreted only on monomer-rich medium and mainly by rhizosphere bacteria and included cellobiohydrolase, glucan endo-1,3-β-glucosidase, lysozyme, α-galactosidase, α,α-trehalase, mannosyl-glycoprotein endo-β-N-acetylglucosaminidase, peptidoglycan lytic transglycosylase and CBMs that bind glucan, maltose (Table S3). One family GH43 with β-xylosidase activity has been identified on polymers media only (xylan and root litter) and secreted by bulk soil bacterium.

### Bulk soil bacteria produce significantly large number of polymer degrading enzymes with signal peptides than rhizosphere bacteria

Microorganisms decompose plant polymers outside the cell and transport the products across the cell wall and membrane as smaller units. As bacteria transfer enzymes across their cellular membranes the amino (N)-terminal signal peptides of their extracellular enzymes facilitate this transfer. We used the conserved amino acid sequences from the signal peptides to predict their presence in extracellular proteins. Bulk soil isolates produced far larger number of GH and CBM proteins with signal peptides than rhizosphere isolates, particularly on plant polymers (Table S4). A number of GHs and CBMs with signal peptides represented a small fraction of total extracellular protein pool detected in rhizosphere bacteria (0.2%), whereas GHs and CBMs with signal peptides were more than twenty times higher in bulk soil bacteria exoproteome (5%) (*P*=0.0001) (Fig. S6A).

Fifteen percent of CAZy proteins were GHs and CBMs with signal peptides produced by rhizosphere bacteria on monomer-rich medium as compared to 41% on average produced by bulk soil bacteria on the same medium (*P*=0.037) (Fig S6B). Less than 5% on average of GHs and CBMs with signal peptides were secreted by the rhizosphere isolates grown with root litter, xylan or cellulose, as opposed to large proportion (72% on average) of enzymes with signal peptides secreted by bulk soil isolates (*P_root litter_* =8.81E-06, *P_xylan_*=1.78E-05, *P_cellulose_*=0.001) (Fig S6B).

### Polymer hydrolysis by bulk soil bacteria associated with production of specific actively secreted GHs and CBMs

To determine the specific enzymes that enable bulk soil bacteria to deconstruct plant polymers and, we identified GHs and CBMs that were significantly more abundant on plant polymeric media (xylan, cellulose and root litter) than on the monomer-rich medium (Fig. 6 A, B, C). GHs and CBMs that were produced at significantly higher amounts on plant polymer media all originated from bulk soil bacteria. These enzymes included 11 glycoside hydrolases (GH2, GH3, GH13, GH15, GH23, GH39, GH43, GH78, GH97, GH106 and GH139) and two carbohydrate binding modules (CBM6 and CBM67) (Fig. 6 A, B, C). They represented a substantial proportion (up to 15%) of the extracellular proteins produced on the plant polymer media and notably most of these GHs and CBMs contained signal peptides (Fig 6 A, B, C). Conversely, GHs and CBMs found to be more abundant in the monomer-rich cultures were produced by both, rhizosphere bacteria (less than 0.6%) and by bulk soil bacteria (less than 3%) (Fig. 6D).

**FIG 6.**
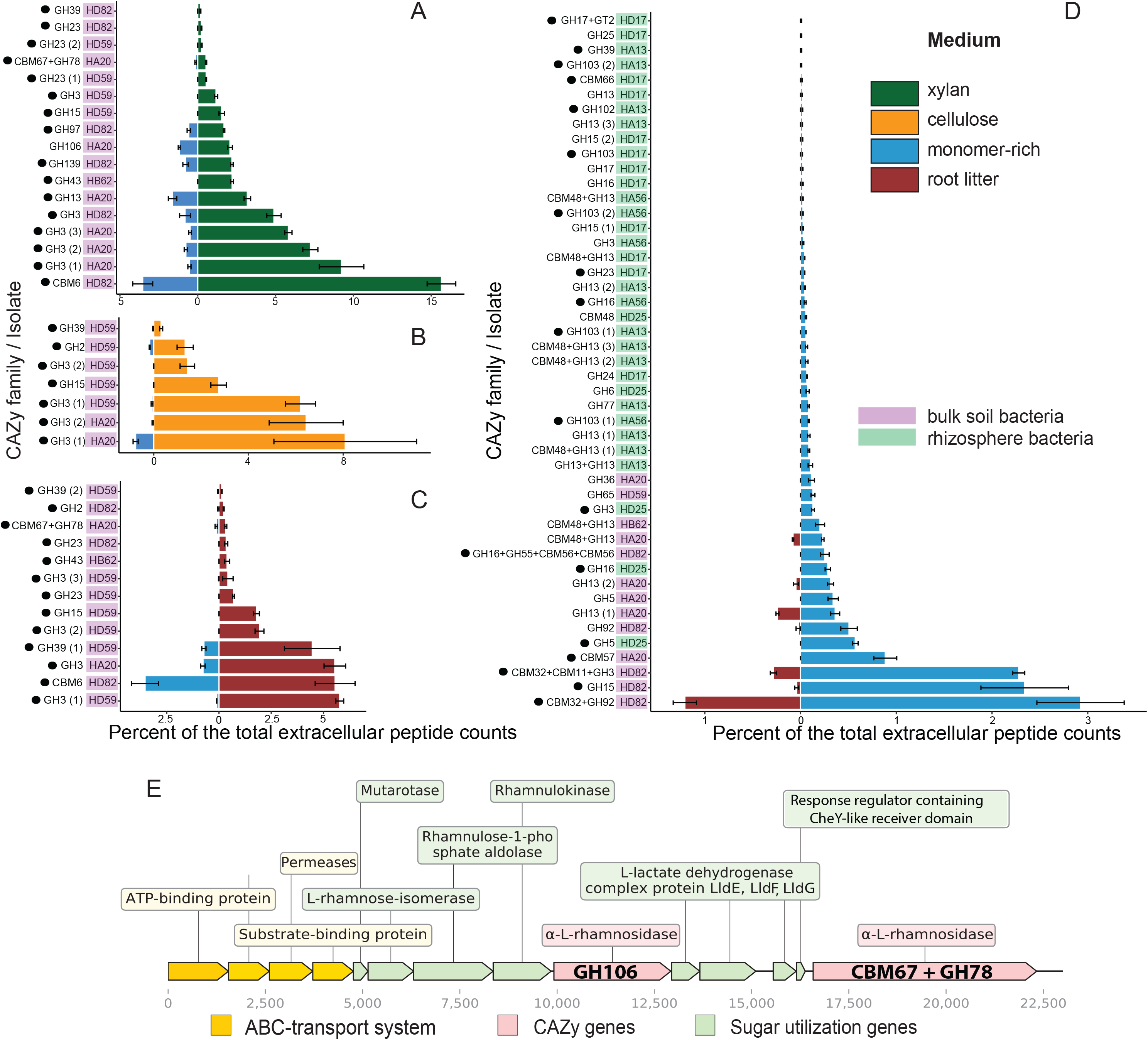
Secreted GHs and CBMs with significantly different abundance on specific medium types (A) xylan, (B) cellulose, (C) root litter, and (D) monomer-rich medium. Significant differences in protein relative abundance across plant polymer- and monomer-rich medium was determined using ANOVA and post hoc Duncan multiple range tests. Each bar graph shows abundance of unique GHs and CBMs produced by each isolate that were significantly higher (*P*<0.05) on any plant polymer medium as opposed to monomer-rich medium (**A**, **B**, **C**) or GHs and CBMs that were higher in monomer-rich medium as opposed to any plan polymer media (**D**). Dots to the left of each protein family ID represent the presence of signal peptides. GHs and CBMs released by bulk soil bacteria vs rhizosphere bacterial isolates are represented by purple or green shading respectively. Error bars represent standard error (n=3). (**E**) Rhamnose transport and metabolism locus of *Nocardioides HA20* showing organization of genes involved in transport and utilization of rhamnose. Each arrow represents a gene in the locus.

Analysis of genomic neighborhoods showed that a subset of the GHs and CBMs we identified occur together in fused domains (e.g. CBM67+GH78) (Fig. 6E), or in CAZyme gene clusters (Fig. S7, Fig 6E), that include at least one CAZyme, one transporter and one transcription factor gene and a number of non-signature genes. A subset of these gene clusters were organized in large polymer utilization loci consisting of co-localized and co-regulated genes involved in sensing, uptake and transformation of polymers (e.g. rhamnose locus, Fig. 6E)(41).

## DISCUSSION

The identification and interpretation of organismal traits and their resulting phenotypic characteristics is key to connecting the functional complexities of organisms to ecosystem function (42, 43). Trait-based approaches can be used to investigate drivers of community assembly and biogeographical patterns, relate microbial diversity and composition to ecosystem function, and predict the consequences of ecosystem perturbation. However, quantifying traits for soil microorganisms can be particularly challenging due to the complexity and diversity of the soil microbiome. New approaches for genome reconstruction from complex communities now permit both the partitioning of organisms into genome-scale response groups and the inference of which genomic features and traits may underlie those responses. To test the resulting hypotheses, phenotyping assays using representative organisms are key to providing functional evidence of genotype to phenotype relationships.

Here, using a collection of bacteria previously classified into response groups based on their growth dynamics in the rhizosphere of a common annual grass, *Avena barbata* and in bulk soil, we derived genomic predictions and provided experimental measurements of metabolic traits hypothesized to contribute to fitness in a bulk versus rhizosphere soil environment. We suggest that the identified traits may explain field-observed successional changes of these bacteria in response to the seasonal C dynamics in a California annual grassland. In this grassland, 40% of photosynthate C is allocated belowground (44), where 20% is lost via exudation and 20% is incorporated in root biomass (44–46). The C exudation is the highest during active plant growth in February–March during active plant growth and declined when grasses began to senesce in late spring (April) (47). In this grassland, most of the detrital roots from the previous year are decomposed by microorganisms and disappear in less than a two-year cycle (44, 47). This C turnover is driven by soil microbial communities, which demonstrate the microbial community succession in this system based on the seasonal availability of different forms of C(10, 20).

Rhizosphere during active plant growth is filled with freshly-released monomeric C molecules exuded by plant roots, including sugars, amino acids, organic acids, nucleotides and nucleosides. Conversely, the bulk soil environment is characterized by an absence of living roots, presence of decomposing recalcitrant highly polymeric organics from litter, and thus is considered resource-limited. Much of the low molecular weight resources are consumed by organisms that have a substrate preference for such compounds (11, 48) or can respond rapidly in the active rhizosphere(4, 49). Following plant senescence, root turnover occurs, facilitated by organisms derived from the bulk soil habitat, adapted to the use of polymers through the costly but necessary production of extracellular enzymes. In this study we demonstrated that bulk soil bacteria have genomic traits that are well aligned with the increased availability of polymeric C and decreased levels of exuded simple C molecules. The genomes of bulk soil bacteria more equipped with polymer-degrading CAZy genes represented by carbohydrate esterases (CE), glycoside hydrolases (GHs), and carbohydrate binding modules (CBMs) compared to the rhizosphere bacteria suggest a greater genomic potential of bulk soil bacteria to utilize polymeric C. Enriched GHs families in the genomes of this group were linked to the potential decomposition of hemicellulose and oligosaccharides. These results suggest that bulk soil bacteria may have a potential fitness advantage encoded in their genomes where plant polymers, particularly hemicellulose, could serve as the main C source when simple C is not available anymore. To confirm the enhanced genomic potential to utilize plant polymers we measured the activity of polymer degrading enzymes in this group of microorganisms and identified that bulk soil bacteria have higher glucosidase and xylosidase activities compared to rhizosphere bacteria when grown on plant polymers. This finding also provides evidence that this group of microorganisms can more effectively utilize plant polymeric C than rhizosphere bacteria. Bulk soil bacteria not only had increased enzymatic activities but also, produced significantly more extracellular enzymes from CAZy families involved in polymer decomposition than rhizosphere bacteria, particularly on plant polymers media. This again provides strong evidence for niche separation based on substrate usage in the rhizosphere versus bulk soil environment and adaptation of bulk soil bacteria to survive in the environment where they have to decompose plant polymers to mine for organic C and N.

Secretion of polymer degrading enzymes is another metabolic trait of bulk soil bacteria analyzed in this study. Bulk soil bacteria were observed to produce more GH and CBM proteins with signal peptides – signal peptides work together with secretory machinery and determine the capability and efficiency of enzyme translocation across the bacterial cytoplasmic membrane (50, 51). Efficient translocation of proteins due to the higher number of signal peptides identified in polymer degrading enzymes of bulk soil bacteria are directly connected to the higher yields of enzymes outside the cell and their extracellular activities.

Finally, identified structural organization of functional genes and different accessory proteins, transcription factors, and transporters into fused domains and clusters associated with degradation of plant polymers by bulk soil bacteria suggest another metabolic trait leading to improved fitness of this group in bulk soil environment. Co-localization and co-transcription of genes involved in the polymer utilization may constitute a fitness advantage for the bacteria due to the cost reduction of the production of protein complexes (52). Such genomic organization synchronizes protein production in protein complexes and helps maintain appropriate protein stoichiometry (52). In addition, fused and co-transcribed GHs and CBM domains can impact enzyme properties including reaction rates and affinity (e.g. see (53)) particularly under high water content conditions (54), in addition to enhancing enzyme thermal stability (e.g. see (55)). Particularly prominent were the CBM6 and CBM67 families produced by bulk soil bacteria on plant polymers. CBM6 is known to have various substrate specificities, including cellulose, xylan, mannan (56), while CBM67 has been reported to bind terminal L-rhamnose sugars and enhance the activity of the associated GH78 α-L-rhamnosidase enzyme (57) These metabolic traits potentially provide another competitive advantage to organisms through enhanced efficiency of polymer deconstruction by the GH enzymes. Utilization of plant polymers represents a biochemically costly strategy and a trait of bulk soil bacteria identified in this study and related to the organization of polymer degrading genes into fused domains and clusters may enhance the efficiency of extracellular enzyme-mediated substrate acquisition in bulk soil bacteria.

The potential for complex niche partitioning and successional shifts in the soil community in response to the plant growth and availability of different forms of C in this annual system has been recently demonstrated using a metatranscriptomic approach (21). Nuccio et al, (2021) revealed multiple microbial ‘guilds’ with a CAZy expression profile specific to a niche defined by both living roots and root detritus, suggesting that a strategy of both root exudate utilization and plant polymer utilization exists. Although we measured traits for two distinct groups of soil bacteria in this study, it is worth noting that this is unlikely a binary scenario and rhizosphere and bulk soil groups of bacteria could co-exist, compete or even share resources. Further work examining metabolic traits of soil microorganisms based on their substrate preferences, resource sharing and trophic interactions should illuminate the fitness of organisms within a diversity of niches present in soil.

### Conclusions

Interactions between plants, soil and microorganisms are dynamic and complex, impacting globally significant biogeochemical processes, such as carbon turnover and stabilization. The metabolic traits of microorganisms underpin their fitness and patterns of community succession. A picture is now emerging of distinct lifestyles of soil microorganisms and their genomic signatures. Amongst numerous defining traits, rhizosphere-adapted bacteria appear to be mostly specialists for low-molecular weight substrates abundant in root exudates. Here we have identified metabolic traits of less-well-studied organisms that are adapted to life in the bulk soil or detritusphere, including heavy investment in plant polymer degrading enzymes and optimization of structural organization of genes associated with polymer degrading potential of bulk soil bacteria. Together, these contrasting lifestyles contribute to the observed succession of soil microbial communities during plant growth. Of course, these substrate preference traits do not occur in isolation, and deciphering how these aspects of microbial metabolism interact with traits related to energy generation, stress response, and interspecies interactions in their natural environment will be important to better understand soil biogeochemical processes and predict the consequences of ecosystem perturbation.

## MATERIALS AND METHODS

### Bacterial isolates and their abundances in soil

Bacterial isolates used in this study (Fig. S1 and Table S1) were isolated from an annual grassland soil at the University of California Hopland Research and Extension Center (HREC) (Hopland, CA, USA; 38° 59’34.5768” N, 123° 4’ 3.7704” W) using media with different nutrient sources and concentrations (11).The abundances of these bacteria in HREC bulk and rhizosphere soil were analyzed previously using 16S rRNA gene sequences mapped to the total community operational taxonomic units (OTUs) identified in this soil (10, 11). Based on the changes in 16S rRNA abundance during the main developmental stages of the dominant annual grass *Avena*, we defined two major groups of bacteria (11). For this study we selected isolates (n=28) with significant change (PERMANOVA, *P*<0.05) in response to plant development identified previously (11). Taxa that significantly increased in abundance in rhizosphere soil while *Avena* was growing are termed ‘rhizosphere bacteria’ (n=19), while taxa that were higher in bulk soil and decreased in abundance in response to plant growth are termed ‘bulk soil bacteria’ (n=9) (Fig. S1). The genomes of isolates used in this study are available in the IMG database (https://img.jgi.doe.gov); accession numbers are provided in Table S1.From 28 isolates, eight were selected for the experiments to measure exoproteome and enzyme activities on media with different C sources (Table S1).

### Analysis of carbohydrate active enzymes in the genomes of bacterial isolates

We predicted the gene copy numbers of carbohydrate active enzymes (CAZy) using a hidden Markov model (HMM) search of protein sequences against the dbCAN HMM database (58–60). The distribution of different CAZy families, fused domains and CAZyme gene clusters were identified (Table S2) using dbCAN2 (58) and their abundance has been normalized by the genome size of an isolate. Signal peptides were identified using SignalP 4.0 (61). Additional screening of polysaccharide utilization loci in the genomes was performed using the IMG portal (https://img.jgi.doe.gov).

### Cultivation of bacterial isolates on plant polymers and monomer-rich media

Eight taxonomically diverse isolates that represent bulk soil bacteria (n=4) and rhizosphere bacteria (n=4) were further selected for the experiment and exoproteome analysis (Table S1). Isolates were grown on four media that represent different sources and forms of carbon (C): (i) monomer-rich medium: ten times diluted R2A medium was used to simulate monomeric labile C sources; (ii) root litter medium: M9 medium with 0.1% ground *Avena* roots supplemented with trace elements and vitamins (62) was used to represent a complex mix of plant polymers; (iii) xylan medium: M9 medium with 0.05% xylan supplemented with trace elements and vitamins was used to represent one of the major plant polymers present in soil; (iv) cellulose medium: M9 medium with 0.1% cellulose supplemented with trace elements and vitamins was used to represent the most abundant plant polymer in the soil environment. Biological triplicates with 35 ml of each media were inoculated with 200 ul of single isolate inoculant (initial OD_600_ = 0.3) and incubated at 28 °C with shaking at 200 r.p.m. for 20 days. Non-inoculated media was used as a negative control. Aliquots of 1.5 ml were used to quantify metabolically active cells using an ATP-bioluminescence assay (CellTiter-Glo® Reagent) and to measure glycosidase and xylosidase activities of the microbial cultures using Matrix Assisted Laser Desorption Ionization (MALDI). For each inoculation, the supernatant was collected by centrifuging at 3300 g for 15 min at 4 °C and then frozen at −80 °C for exoproteome analysis.

### Polymer-degrading enzymatic activity analyses using Matrix Assisted Laser Desorption Ionization Mass Spectrometry (MALDI-MS)

For enzyme activity analyses, we used fluorous-tagged substrates (Cellobiose-F17: chemical formula C_36_H_52_F_17_N_5_O_14_, exact mass 1101.3239 and Xylobiose-F17: chemical formula C_34_H_48_F_17_N_5_O_12_, exact mass 1041.30) (63). 9 μl of sample was mixed with 1 μl of each substrate and incubated at 28 °C with shaking at 200 r.p.m. for 3 h. After this incubation, 10 μl of ethanol was added to each sample to achieve a final substrate concentration of 25 μM. All samples were diluted 1:2 with 90/10 water/ethanol containing an internal standard with a final concentration of 3.33 uM. Then, to 4 μL of the diluted samples, 4 μL universal MALDI matrix (20 mg/mL-1 in 1:3 (v/v) H_2_O:MeOH; Sigma-Aldrich) was added. Samples were printed in four technical replicates onto a stainless-steel blank MALDI plate using an ATS-100 acoustic transfer system (BioSera) with a sample deposition volume of 10 nl. Samples were printed in clusters of four replicates, with the microarray spot pitch (center-to-center distance) set at 900 μm. MS-based imaging was performed using an ABI/Sciex 5800 MALDI TOF/TOF mass spectrometer with laser intensity of 3,500 over a mass range of 750–2,000 Da. Each position accumulated 20 laser shots. The instrument was controlled using the MALDI-MSI 4800 Imaging Tool. Surface rasterization was oversampled using a 75 μm step size. Average signal intensities for the ions of fluorous-tagged substrates per sample spot were determined using the OpenMSI Arrayed Analysis Toolkit (64). Negative controls of non-enzymatic hydrolysis were subtracted to correct the calculated activities; the signal intensity of the internal standard was used for normalization.

#### Exoproteome extraction

For exoproteome analysis, each sample was thawed, and 15 ml transferred to chemically compatible polypropylene 50 ml tubes (Olympus Plastics, Waltham, MA). 35 ml of ice-cold 2:1 (v/v) chloroform:methanol was added, vortexed and centrifuged at 4500 g for 10 min at 4 °C to separate proteins. The upper and lower phases were removed and the protein interphase was washed with 10 ml of ice-cold methanol and re-centrifuged to pellet the protein. Ice-cold methanol (1ml) was added again to the protein pellet and transferred into 1.5 ml tubes centrifuged at 12000 g for 10 min at 4 °C. The methanol was decanted and the protein pellet was allowed to dry inverted on a Kim wipe for 20 min. To each protein pellet, 100 μl of UPX Universal Protein Extraction buffer (Expedeon, San Diego, CA) was added and water-bath sonicated into solution. Each sample was incubated at 95 °C for 5 min to ensure reduction and solubilization of protein. The samples were then vortexed and sonicated for 2 min, lightly spun to collect the condensate and allowed to cool at 4 °C for 45 min. The samples were then centrifuged at 15000 g for 10 min and Filter Aided Sample Preparation (FASP) (65) kits were used for protein digestion (Expedeon, San Diego, CA) according to the manufacturer’s instructions. Briefly, 400 μl of 8 M urea (all reagents included in the kit) was added to each 500 μl 30K molecular weight cut off (MWCO) FASP spin column and up to 100 ul of the sample in UPX buffer was added, centrifuged at 14000 g for 30 min to bring the sample to the dead volume. The waste was removed from the bottom of the tube and another 400 μl of 8 M urea was added to the column and centrifuged again at 14000 g for 30 min and repeated once more. 400 μl of 50 mM ammonium bicarbonate (provided) was added to each column and twice centrifuged for 20 min. The column was placed into a new fresh, clean and labeled collection tube. Digestion solution was made by dissolving 4 μg trypsin in 75 μL 50 mM ammonium bicarbonate solution and added to the sample. Each sample was incubated for 3 h at 37 °C with 800 r.p.m. shaking on a thermomixer with a thermotop (Eppendorf, Hamburg, Germany) to reduce condensation into the cap. The resultant peptides had 40 μl of ammonium bicarbonate solution added and were then centrifuged through the filter and into the collection tube at 14000 g for 15 min. 40 μl of ammonium bicarbonate solution was added to the filter, which was then centrifuged again. Peptides were concentrated to 30 μL using a SpeedVac. Final peptide concentrations were determined using a bicinchoninic acid (BCA) assay (Thermo Scientific, Waltham, MA USA). Each sample was diluted to 0.1μg/μl and vialed for mass spectrometry analysis.

#### Liquid chromatography mass spectrometry analysis of extracted exoproteome

LC-MS/MS analysis for each sample was performed using a ThermoFinnigan Q-Exactive ion trap mass spectrometer at Pacific Northwest National Laboratory, Richland WA, outfitted with a custom-built electrospray ionization (ESI) interface. Samples were loaded onto a 70-cm Phenomenex Jupiter C18 column (3 μm diameter porous, inner diameter: 75 μm). Data were acquired for 100 min, beginning 15 min into the gradient. Precursor MS spectra (AGC 3 × 106) were collected over 400–2000 m/z followed by data-dependent MS/MS spectra (AGC 1 × 105) of the twelve most abundant ions, using a collision energy of 30 %. A dynamic exclusion time of 30 s was used to discriminate against previously analyzed ions. Mobile phase A was 0.1 % v/v formic acid in water and mobile phase B was 0.1 % v/v formic acid in acetonitrile. The gradient was generated using a Waters nanoACQUITY UPLC constant flow LC system.

#### Identification and functional screening of exoproteomes

For peptide identification, mass spectra from the resulting analyses were evaluated using the MSGF+ software (66) and searched against protein-coding open reading frames of eight isolate genomes used in this experiment. The predicted open reading frame (e.g., predicted protein) libraries were created using both the forward and reverse direction to allow determination of a False Discovery Rate (FDR). MSGF+ was then used to search the experimental mass spectra data against both the forward/reverse decoy databases. Data were filtered by MSGF+ spectra probability (<1 × 10^10^, equivalent to a BLAST evalue), mass accuracy (-/+ 5 ppm), protein level FDR of 1% and minimum of two unique peptides per protein identification. Identified peptide must be present in two out of three replicates of the same treatment. The relative abundance of CAZy, GHs, CBMs represented as a percentage of the total extracellular peptide counts identified in 0.1μg/μl of analyzed peptides extracted from each sample (Table S2). All mass spectrometry proteomics data from this study have been deposited in the MassIVE repository (https://massive.ucsd.edu) with the accession number MSV000084044.

#### Statistical analyses

All statistical analyses were performed within the R software environment (R Development Core Team, 2017) using the ‘vegan’ and ‘agricolae’ packages (67) (68). Kruskal–Wallis one-way analysis of variance was used to test for statistical differences in the distribution of carbohydrate active enzymes present in the genomes of bulk soil and rhizosphere bacteria, and to test for differences of produced extracellular CAZy between these two groups of bacteria. Analysis of variance and post-hoc Duncan’s multiple range test were also used to test for significant differences in the composition of glycoside hydrolases and carbohydrate binding modules released by bulk soil and rhizosphere bacteria across the different carbon sources (monomer-rich, root litter, xylan and cellulose media).

## Acknowledgements

This study was supported by the DOE, Office of Science, Office of Biological Environmental Research, including Genomic Sciences program awards no. DE-SC0010570, DE-SC0016247 and DE-SC0014079 to MKF and awards SCW1589 and SCW1678 to JPR. Isolate genome sequencing was conducted by the U.S. Department of Energy Joint Genome Institute, a DOE Office of Science User Facility, under a Community Science Program award to ELB, supported by the Office of Science of the U.S. Department of Energy under Contract No. DE-AC02-05CH11231. Work performed at the Lawrence Berkeley National Laboratory including DOE Early Career Award to TRN was carried out under contract DE-AC02-05CH11231. This work in part was performed in the Environmental Molecular Sciences Laboratory, a national scientific user facility sponsored by the Department of Energy (DOE) Office of Biological and Environmental Research, and located at Pacific Northwest National Laboratory (PNNL). PNNL is a multi-program national laboratory operated by Battelle for the DOE under Contract DE-AC05-76RLO 1830. Work conducted at Lawrence Livermore National Laboratory was supported under the auspices of the U.S. DOE under Contract DE-AC52-07NA27344.

## Author contributions

K.Z., T.R.N., M.K.F., J.P.R., M.S.L. and E.L.B. developed the hypotheses. K.Z., C.D.N., M.d.R. and K.D. performed the experimental analyses. K.Z., R.A.W., U.K. and analyzed the data. K.Z., T.R.N., M.K.F. and E.L.B. wrote the paper. All authors provided comments and edits on the manuscript.

