## Supplementary Information for "Metabolic traits and the niche of bulk soil bacteria in a Mediterranean grassland"

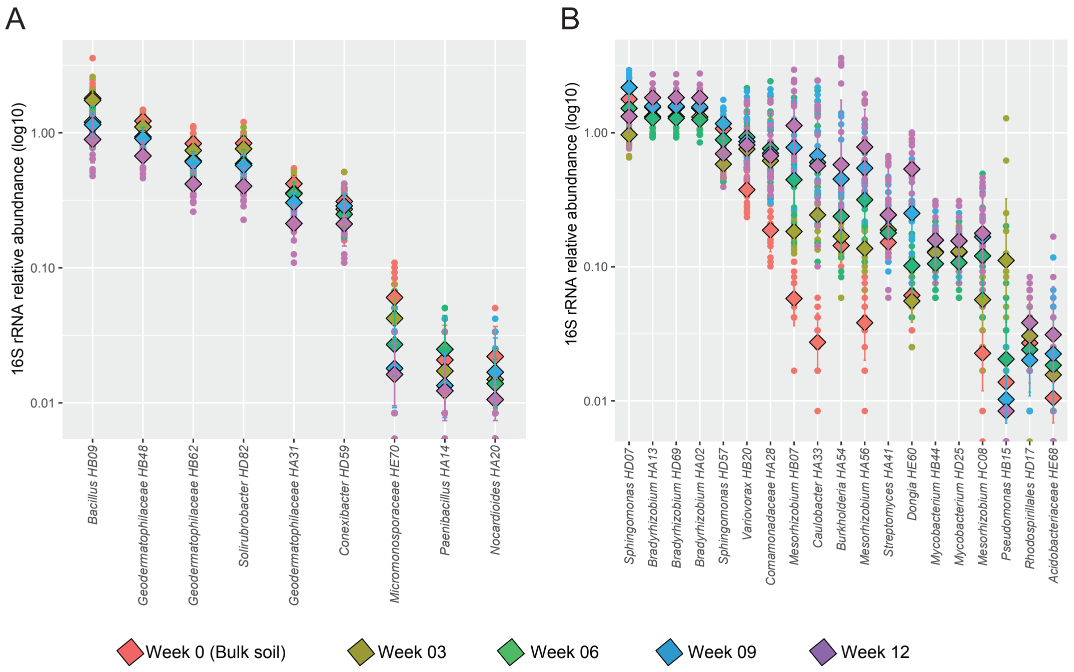


**Fig. S1** Changes in 16S rRNA gene relative abundance of bacterial isolates during main developmental stages of *Avena* growth (Week 0 – Week 12) and in bulk soil. (**A**) Relative abundance of bulk soil bacteria, (**B**) relative abundance of rhizosphere bacteria. In each dot plot a point of the same color denotes a single measurement of 16S rRNA gene abundance of the same isolate. The diamond represents mean of the abundance for each week and error bars show standard deviation. Modified from Zhalnina et al., 2018.

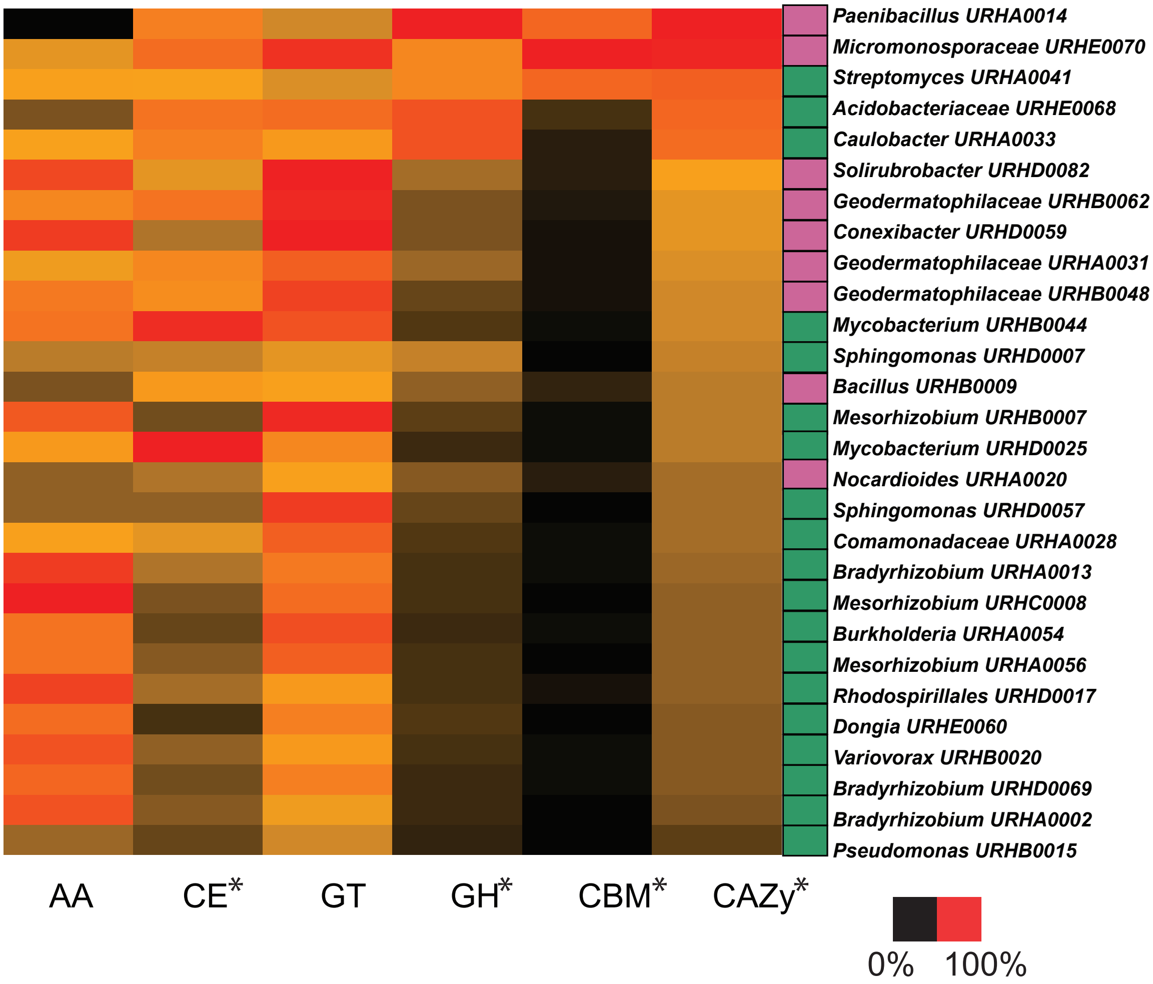


**Fig. S2** Heatmap of genes number in carbohydrate active enzyme (CAZy) families in the genomes of 28 isolates from a California annual grassland soil. These isolates were classified into two groups based on their response to growth of the common annual grass *Avena barbata* in previous studies {zhalnina:2018il}: i) bulk soil bacteria, which were more abundant in bulk soil and declined in relative abundance in response to the plant growth (purple), (ii) rhizosphere bacteria, which increased in abundance during plant growth (green). AA – auxiliary activities enzymes; CE – carbohydrate esterases; GT – glycosyl transferases; GH – glycoside hydrolases; CBM - carbohydrate binding modules. Gene copy number was normalized by the genome size of each isolate. Heatmap shows the abundance of genes within the same family on a scale 0-100% where 100 is the highest number of genes and 0 is the lowest number of genes within the same family across all isolates. Differences in the distributions of traits between the two groups of isolates (rhizosphere and bulk soil bacteria) were evaluated using a Kruskal–Wallis one-way analysis of variance test (*$P<0.05$).

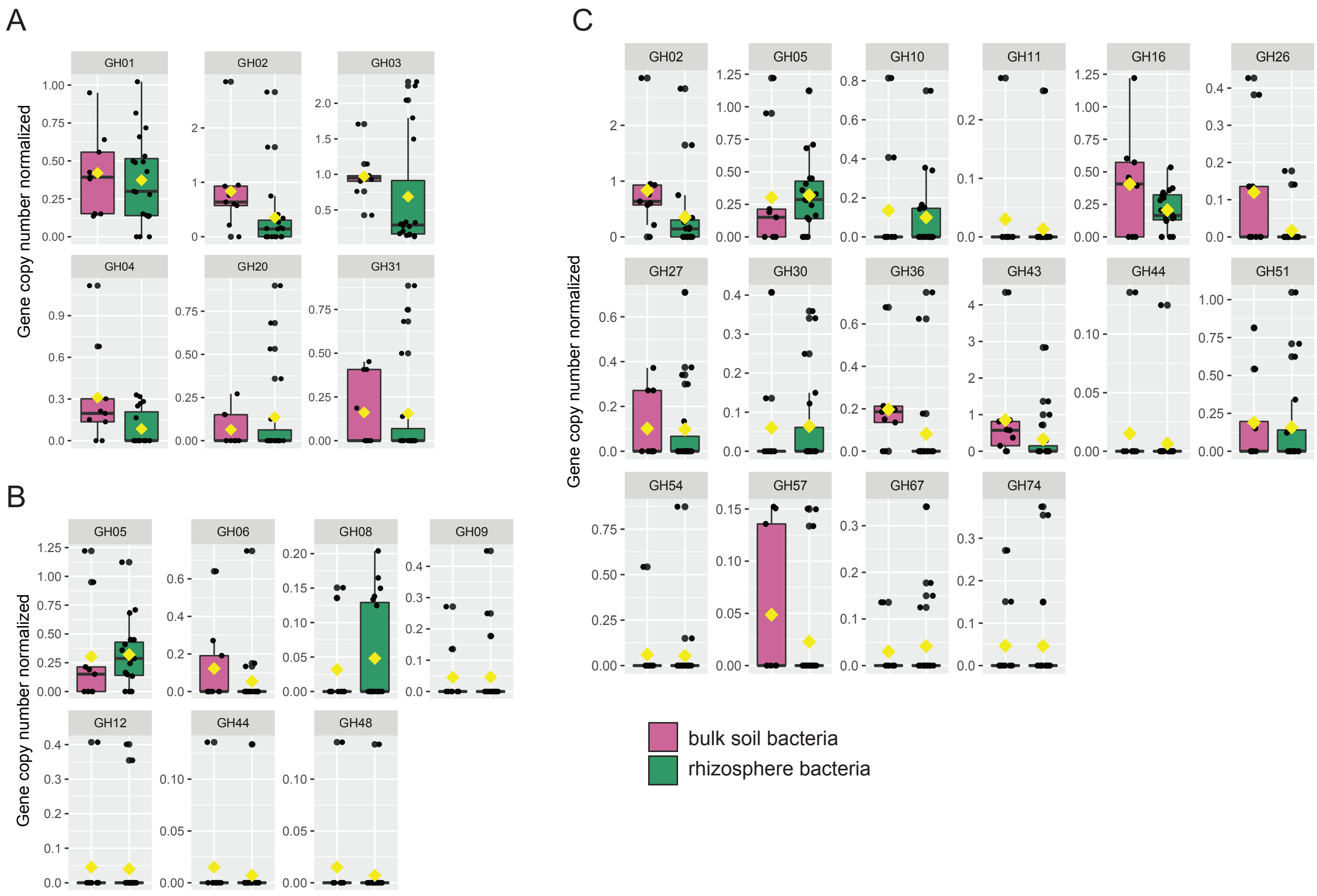


**Fig. S3 Glycoside hydrolase (GH) families involved in decomposition of polymeric substrates of bulk soil and rhizosphere bacteria. (A)** GHs involved in oligosaccharide, (**B**) cellulose and (**C**) hemicellulose deconstruction.

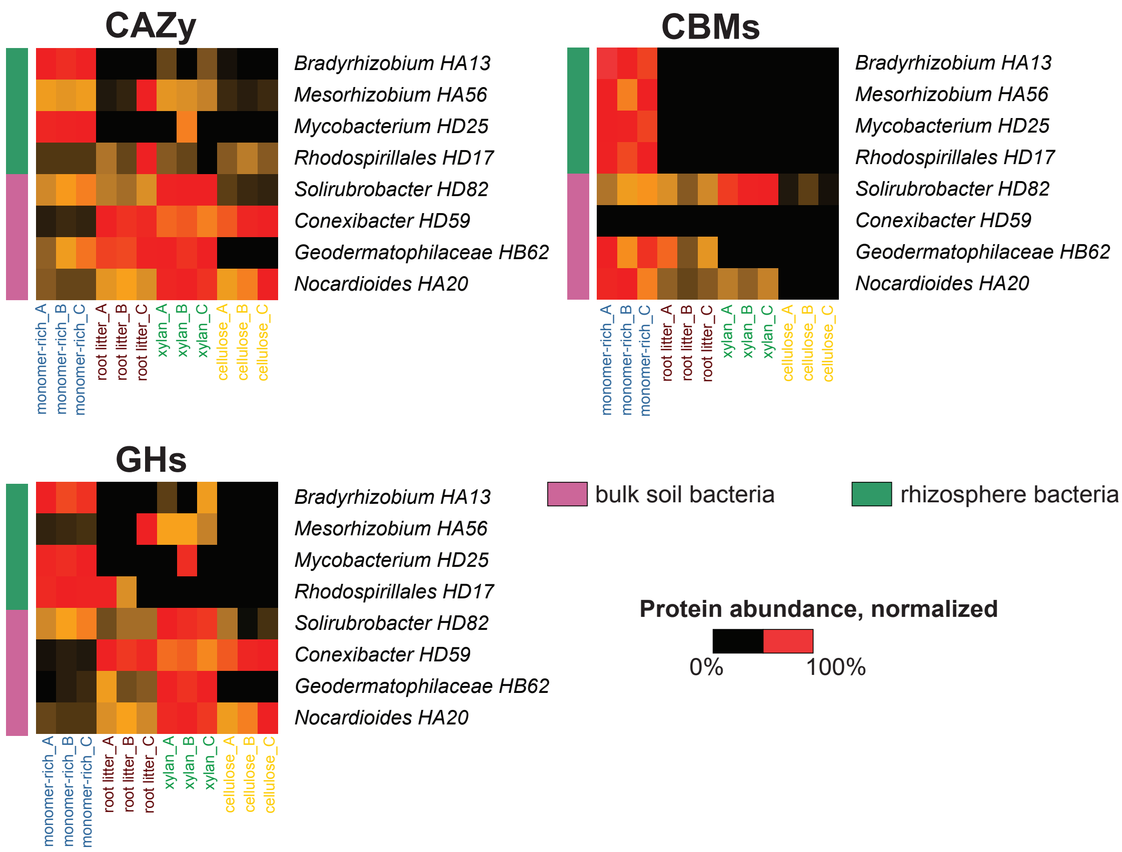


**Fig. S4 CAZy classes of extracellular proteins produced by bulk soil and rhizosphere bacteria on different media.** Four media with different carbon sources (monomer-rich, root litter, xylan and cellulose) were inoculated with bulk soil and rhizosphere isolates. Spent media was collected after 20 days of bacterial growth and analyzed. Protein abundance was scaled to the protein with maximum abundance for each isolate.

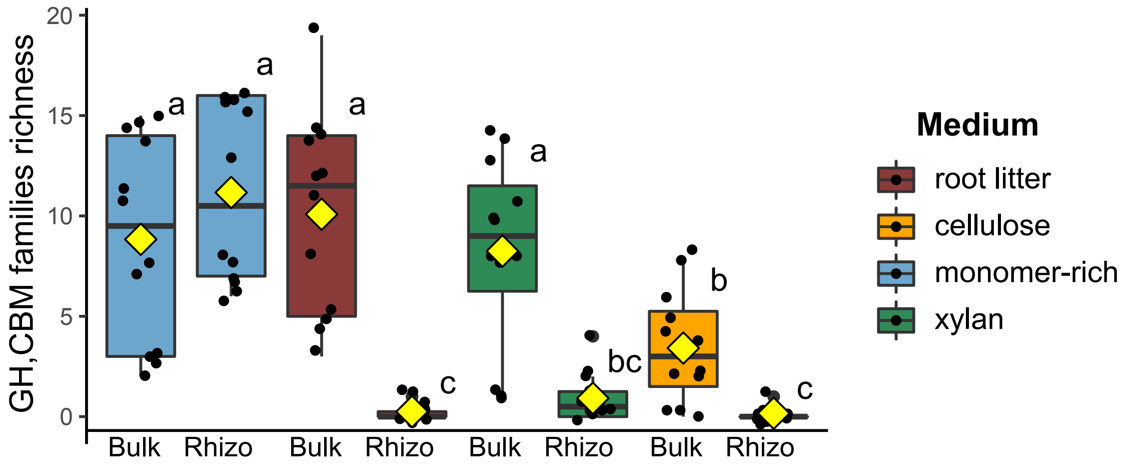


**Fig. S5 Richness of CBM and GH families produced by bulk soil and rhizosphere bacteria on four different media.** Richness of GH and CBM families in bulk soil and in rhizosphere bacteria, represented as a unique GH and CBM families counts identified in each sample. Significant differences in GHs and CBMs richness between treatments was determined using ANOVA and post hoc Duncan multiple range tests. Different letters represent significantly different treatments ($P$< 0.05).

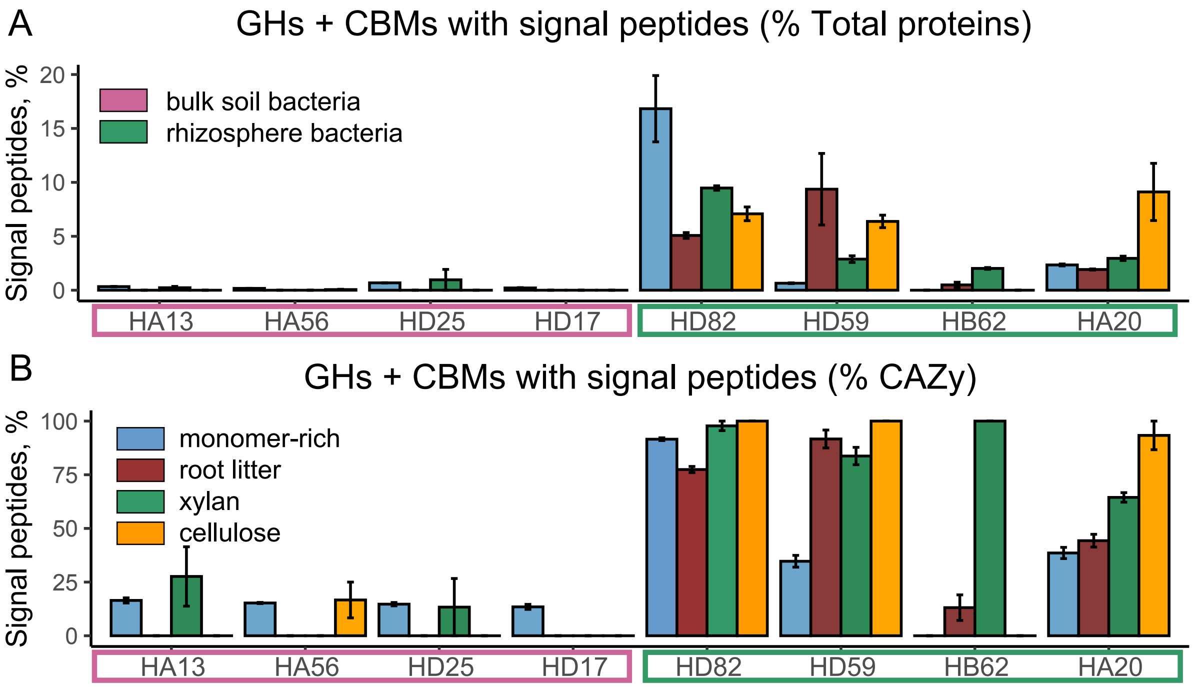


**Fig. S6 CBMs and GHs with signal peptides.** Number of CBMs and GHs represented as a percent of total peptides (**A**) or as a percent of CAZy (**B**) identified on four different media inoculated by bulk soil and rhizosphere bacteria. Number of replicates for each treatment n=3. Error bars represent standard error.

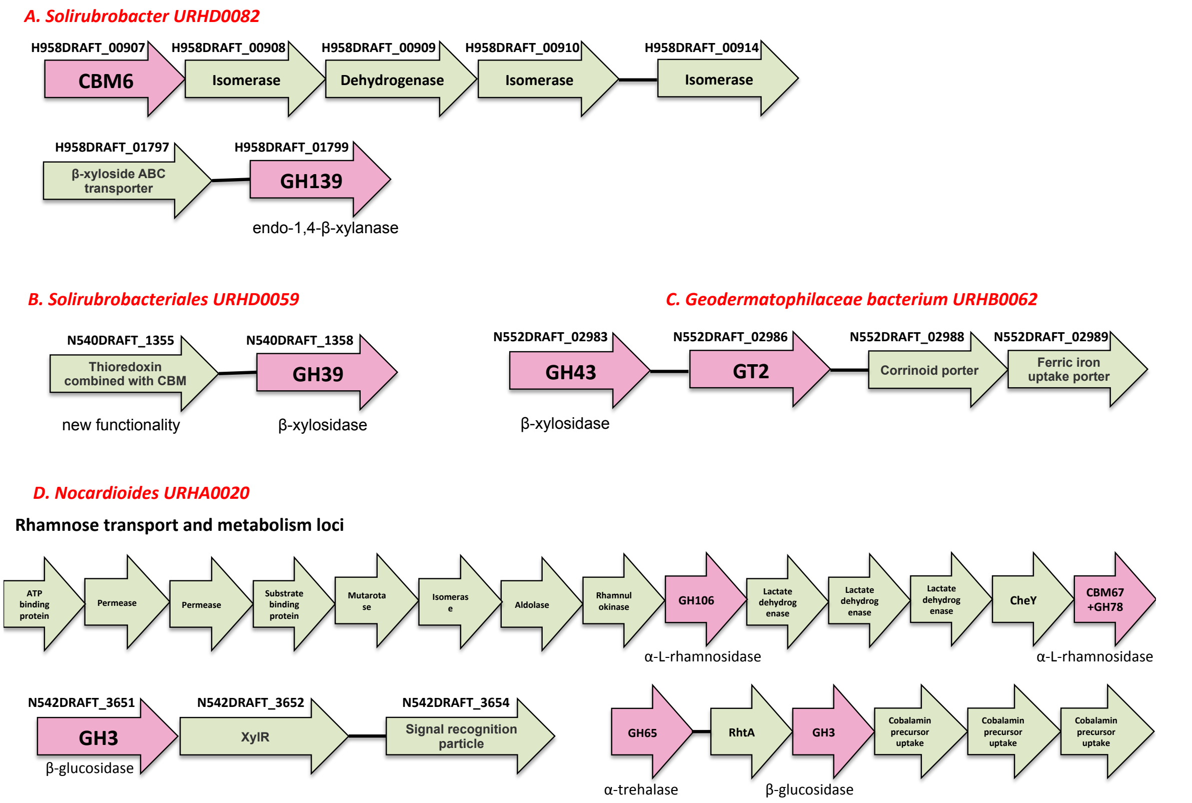


**Fig. S7** CAZyme gene clusters (CGCs) and polymer utilization loci. CGCs must include at least one CAZyme, one transporter and one transcription factor gene and the number of non-signature genes. CAZy genes shown in pink color.
